# Leakiness at the human-animal interface in Southeast Asia and implications for the spread of antibiotic resistance

**DOI:** 10.1101/2021.03.15.435134

**Authors:** Maya L. Nadimpalli, Marc Stegger, Roberto Viau, Vuthy Yith, Agathe de Lauzanne, Nita Sem, Laurence Borand, Bich-tram Huynh, Sylvain Brisse, Virginie Passet, Søren Overballe-Petersen, Maliha Aziz, Malika Gouali, Jan Jacobs, Thong Phe, Bruce A. Hungate, Victor O. Leshyk, Amy J. Pickering, François Gravey, Cindy M. Liu, Timothy J. Johnson, Simon Le Hello, Lance B. Price

## Abstract

International efforts to curb antimicrobial resistance have focused on drug development and limiting unnecessary use. However, in areas where water, sanitation, and hygiene infrastructure is lacking, and where biosecurity in food-animal production is poor, pathogen-flow between humans and animals could exacerbate the emergence and spread of resistant pathogens. Here, we compared mobile resistance elements among *Escherichia coli* recovered from humans and meat in Cambodia, a country with substantial connectivity between humans and animals, unregulated antibiotic use, and poor environmental controls. We identified multiple resistance-encoding plasmids and a novel, *bla*_*CTX-M*_ and *qnrS1*-encoding transposon that were widely dispersed in both humans and animals, a phenomenon rarely observed in high-income settings. Our findings indicate that plugging leaks at human-animal interfaces should be a critical part of addressing antimicrobial resistance in low and middle-income countries.

## Introduction

Low and middle-income countries are projected to experience the greatest mortality and economic fallout from the looming antimicrobial resistance crisis. Global public health organizations are collaborating with countries to develop “One Health”-oriented national action plans aimed at improving antimicrobial stewardship in both humans and animal production.

However, the importance of environmental controls at the human-animal interface has not received the same level of attention as antimicrobial use. This lack of prioritization may be due to high-income country bias, as studies from the United Kingdom (Day et al., 2019) and the European Union (Mughini-Gras et al., 2019) have identified only minimal sharing of antibiotic-resistant bacteria between humans and food-animals. However, the situation could be vastly different in low and middle-income countries where environmental controls at the human-animal interface are lacking (**Fig. 1**).

**Fig. 1.**
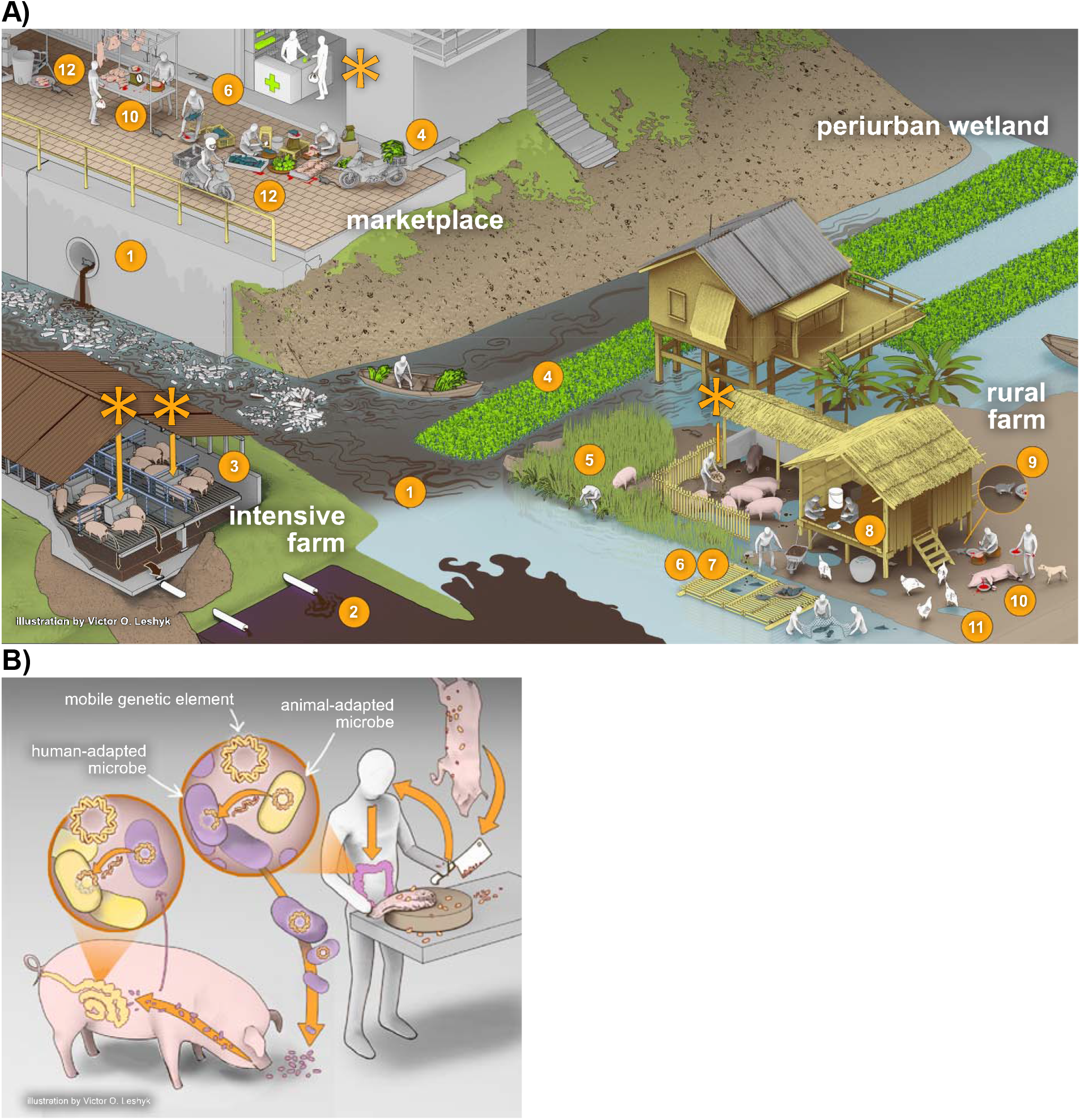
Unchecked environmental leaks and antimicrobial use create myriad opportunities for the exchange of bacteria and mobile resistance elements among humans and animals. Panel A) depicts potential transmission pathways: 1) Lack of sewage treatment infrastructure leads to contamination of drinking water for humans and animals; 2) Waste from concentrated animal feeding operations contaminated drinking water for humans and animals; 3) Raising many animals in confinement propagates disease; 4) Untreated human sewage fertilizing crops for human consumption; 5) Open defecation leads to fecal-oral transmission among humans and animals consuming human feces; 6) Direct contact with unsafely managed animal waste; 7) Farmed fish are fed feces from pigs and poultry; 8) Consumption of undercooked chicken and fish; 9) Vermin and flies contaminate food preparation areas; 10) Meat easily contaminated by feces during slaughtering and processing in urban and rural areas; 11) Multiple animal species in contact in unhygienic conditions; 12) Poor food hygiene in markets. Panel B) depicts how frequent mixing of human- and animal-adapted microbes creates opportunities for the exchange of antibiotic resistance at multiple scales; *i*.*e*., of bacteria, plasmids, and transposable elements. *Note*: Layout stylized to indicate connectivity. Asterisks indicate antibiotic inputs.

Water, sanitation and hygiene (WASH) in human communities and “biosecurity” in animal production aim to control the flow of pathogens in and out of these systems. In settings where humans and food-animals are living in close proximity, the absence or breakdown of these controls - hereafter referred to as “leakiness” – could have profound implications: the continuous, circular exchange of resistant bacteria selected through unregulated antibiotic use could facilitate the exchange of resistance genes among bacteria in both sectors. Here, we investigated this hypothesis by using a multi-scale, molecular approach to explore the spread of mobile resistance elements between humans and animals in Phnom Penh, Cambodia, a highly leaky urban center in Southeast Asia.

East and Southeast Asia are hotspots for emerging zoonotic infectious diseases such as highly pathogenic avian influenza H5N1, severe acute respiratory syndrome (SARS-CoV-1 and 2), and Nipah virus, all of which have emerged in the past few decades. Cambodia is among the poorest countries in Southeast Asia and 76% of the estimated 16 million residents live in rural areas (National Institute of Statistics, Ministry of Planning, 2018). As in other Southeast Asian countries, Cambodian livestock production is rapidly increasing to support growing demand.

Between 2012 and 2016, domestic poultry production increased by 53% to 35.7 million chickens per year and pig production by 34% to 2.9 million pigs per year (Ministry of Agriculture, Forestry, and Fisheries (MAFF), 2017), and meat demand is expected to increase by another 20% by 2024 (Heng Sen, 2018). While commercial livestock production has nearly doubled during the past decade (Ministry of Agriculture, Forestry, and Fisheries (MAFF), 2017), at least 80% of livestock continues to be raised by resource-constrained households located in rural areas (Ministry of Agriculture, Forestry, and Fisheries (MAFF), 2017; National Institute of Statistics, Ministry of Planning, 2018).

WASH conditions and animal husbandry practices in Cambodia suggest ample opportunity for bidirectional exchange of fecal-oral pathogens between humans and livestock (**Fig. 1**). First, Cambodia has the highest rate of open human defecation in the region (UNICEF Cambodia, 2019); 26.2% of rural residents and 7.7% of urban residents outside the capital continue to defecate in fields or other open spaces (National Institute of Statistics, Ministry of Planning, 2018). Because most livestock raised in rural and urban households are allowed to roam freely during the day, these animals may consume human feces. Second, nearly half of livestock-owning households report dumping untreated animal manure directly into the environment (Gunilla Ström et al., 2018) suggesting that untreated drinking water may be a source of animal fecal bacteria. Cambodia has the lowest access to piped drinking water in Southeast Asia, and nearly 1 in 5 urban residents outside the capital and 1 in 2 rural residents cannot reliably access clean water (National Institute of Statistics, Ministry of Planning, 2018). Third, poor WASH practices along the food supply chain in Cambodia, especially in antiquated slaughterhouses and crowded live-animal markets, facilitate the contamination of meat and produce by animal and human fecal bacteria. Finally, behaviors like using untreated animal manure to fertilize crops for human consumption (Gunilla Ström et al., 2018) and eating undercooked chicken and fish (Nadimpalli et al., 2019b) likely exacerbate the spread of zoonotic pathogens. With so many leaks between human communities and animal production systems, antimicrobial use in both sectors could lead to the emergence and spread of antimicrobial-resistant bacteria.

We explored the antimicrobials most commonly used in human medicine and food-animal production in Southeast Asia and found substantial overlap. Antibiotics administered to chickens and pigs included all antibiotics that are commonly used in human medicine, including third-generation cephalosporins, penicillins, fluoroquinolones, gentamicin, and co-trimoxazole (Choisy et al., 2019; G. Ström et al., 2018; Carrique-Mas et al., 2015; Om et al., 2016). Farmers in Cambodia reported purchasing human antimicrobials for their animals when veterinary drugs were ineffective (G. Ström et al., 2018) and antibiotics that are critically important for human medicine (*e*.*g*. colistin) were used extensively in poultry production (Carrique-Mas et al., 2015). Such practices mean that the same antibiotic resistance genes could confer a selective advantage in multiple hosts. Previously, we found that extended-spectrum β-lactamase (ESBL)-producing *Escherichia coli* colonizing healthy community members in Phnom Penh, Cambodia were highly similar to strains recovered from meat and fish sold at markets (Nadimpalli et al., 2019b). For example, one-third of human-origin ESBL-producing *E. coli* strains encoded the *bla*_CTX-M-55_ gene, which was the predominant ESBL gene type among *E. coli* recovered from meat and fish (**Fig. 2a**). This sharply contrasts findings from multiple high-income counties where WASH and biosecurity infrastructure are robust (Day et al., 2019; Mughini-Gras et al., 2019); for example, a recent study from the UK found only 5% of human-origin ESBL-producing *E. coli* strains harbored ESBL genes that predominated among meat (Day et al., 2019) (**Fig. 2a**). We hypothesized that substantial leakiness between humans and animals in Cambodia, combined with similar antibiotic pressures in both reservoirs, could be contributing to the synergistic amplification and mutual exchange of mobile resistance elements.

**Fig. 2.**
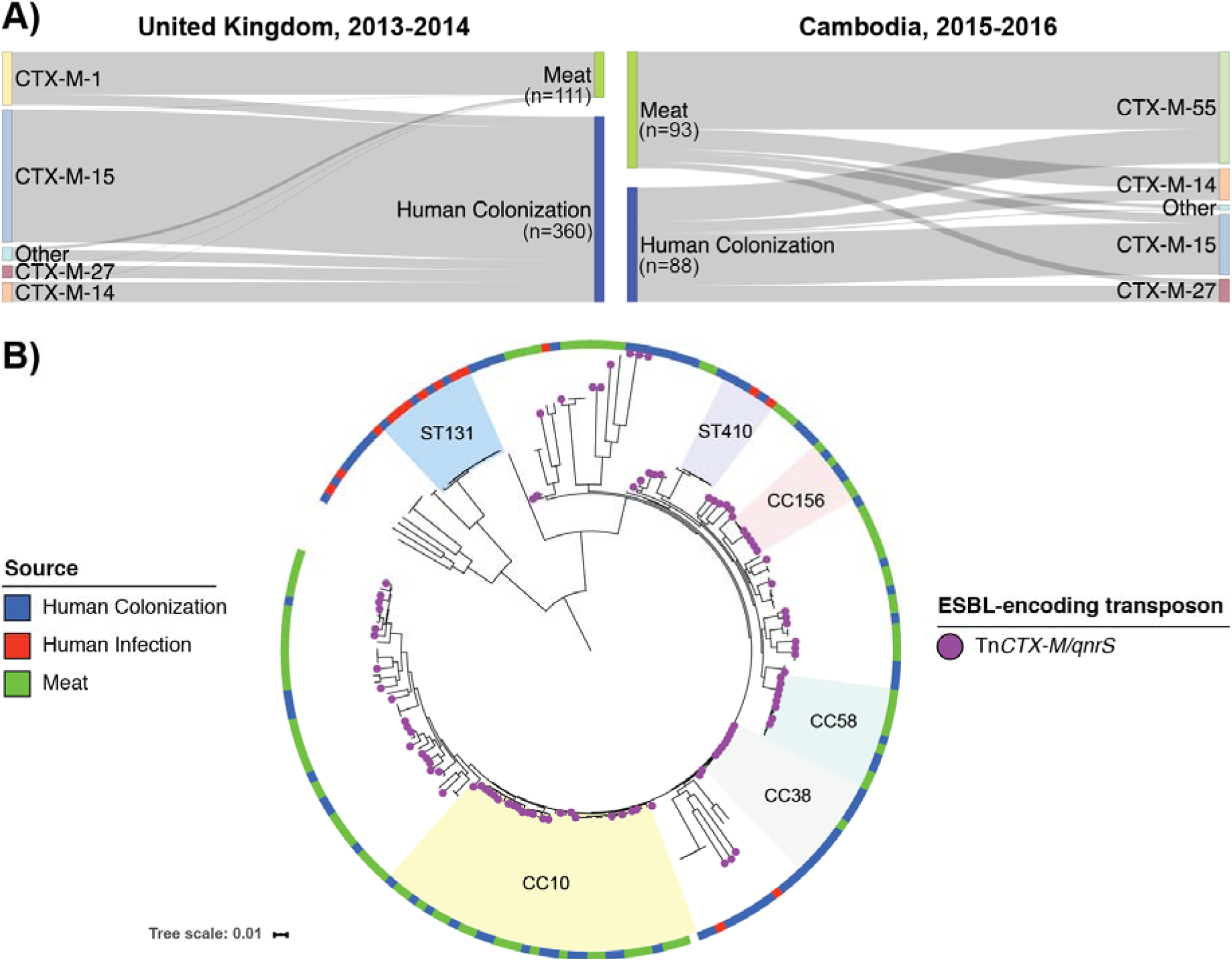
High antibiotic use and environmental leakiness is associated with the mobilization of an ESBL-encoding transposon across different bacterial lineages and hosts in Cambodia. A) Sankey diagram depicting the flow of ESBL genes among *Escherichia coli* from human and animal origin; arrow width is proportional to the flow rate. ESBL genes flow freely among human and animal-origin *E. coli* in the leaky Cambodian ecosystem but are largely isolated to specific hosts in the less leaky UK ecosystem. B) Core-genome phylogenetic tree depicting the evolutionary relationships of ESBL-encoding *E. coli* isolates from Cambodia. A bla_*CTX-M*_-encoding transposon has been acquired by multiple *E. coli* sub-lineages from humans and meat, underscoring rampant genetic exchange in a highly leaky system with substantial antimicrobial selective pressure.

Here, we analyzed ESBL-encoding elements among *E. coli* of human and animal origin to determine if strains from these two sources shared a common pool of mobile resistance elements. We first used long-read sequencing to assemble high-quality draft genomes of five ESBL-producing *E. coli* isolates from the feces of healthy humans (n=2) and from pork meat (n=2) and chicken (n=1) sold at Phnom Penh markets (**Supplementary Text, Tables S1-S2, Fig. S1**). We identified the ESBL-encoding plasmids they harbored, annotated them, and then screened Cambodian collections of ESBL-producing *E. coli* from healthy, gut-colonized humans (n=88), clinical specimens (n=15), and meat and fish from markets (n=93) for their presence (**Data S1**).

## Results and Discussion

We identified four distinct *bla*_CTX-M-55_-encoding plasmids that were shared with slight variations among *E. coli* isolates of both human and animal origin (**Fig. 3**). One IncHI1-type plasmid (pC27A-CTX-M-55) co-encoding resistance to third-generation cephalosporins (*bla*_CTX-M-55_), fluoroquinolones (*qnrS1*), phenicols (*floR*), tetracyclines (*tet*(A)), sulfonamides (*sul3*), trimethoprim (*dfrA14*), and aminoglycosides (multiple genes) was present in *E. coli* from chicken meat (n=3), fish (n=12), pork meat (n=2) and healthy humans (n=2). An IncHI2-type plasmid (pP59A-CTX-M-55) co-encoding resistance to β-lactams (*bla*_CTX-M-55_, *bla*_TEM-1-B_), fluoroquinolones (*qnrS1*), macrolides (*mef*(B)), phenicols (*floR*), sulfonamides (*sul2, sul3)*, and aminoglycosides (*aac(3)-Iid, aad24*) was prevalent among pork meat isolates (n=6) and identified in a healthy human (n=1). An IncF[F18:A-:B1]-type plasmid (pP225M-CTX-M-55) co-encoding resistance to third-generation cephalosporins (*bla*_CTX-M-55_), tetracyclines (*tet*(A)), and aminoglycosides (*aadA1*) was primarily found among isolates from healthy humans (n=6) but was also carried by *E. coli* from a human urinary tract infection (n=1) and from fish (n=1). Finally, a novel plasmid type (p276M-CTX-M-55) co-encoding resistance to third-generation cephalosporins (*bla*_CTX-M-55_), fluoroquinolones (*qnrS1*), phenicols (*floR*), and tetracyclines (*tet*(A)) was identified from a healthy human isolate (n=1) and from chicken meat (n=1).

**Fig. 3.**
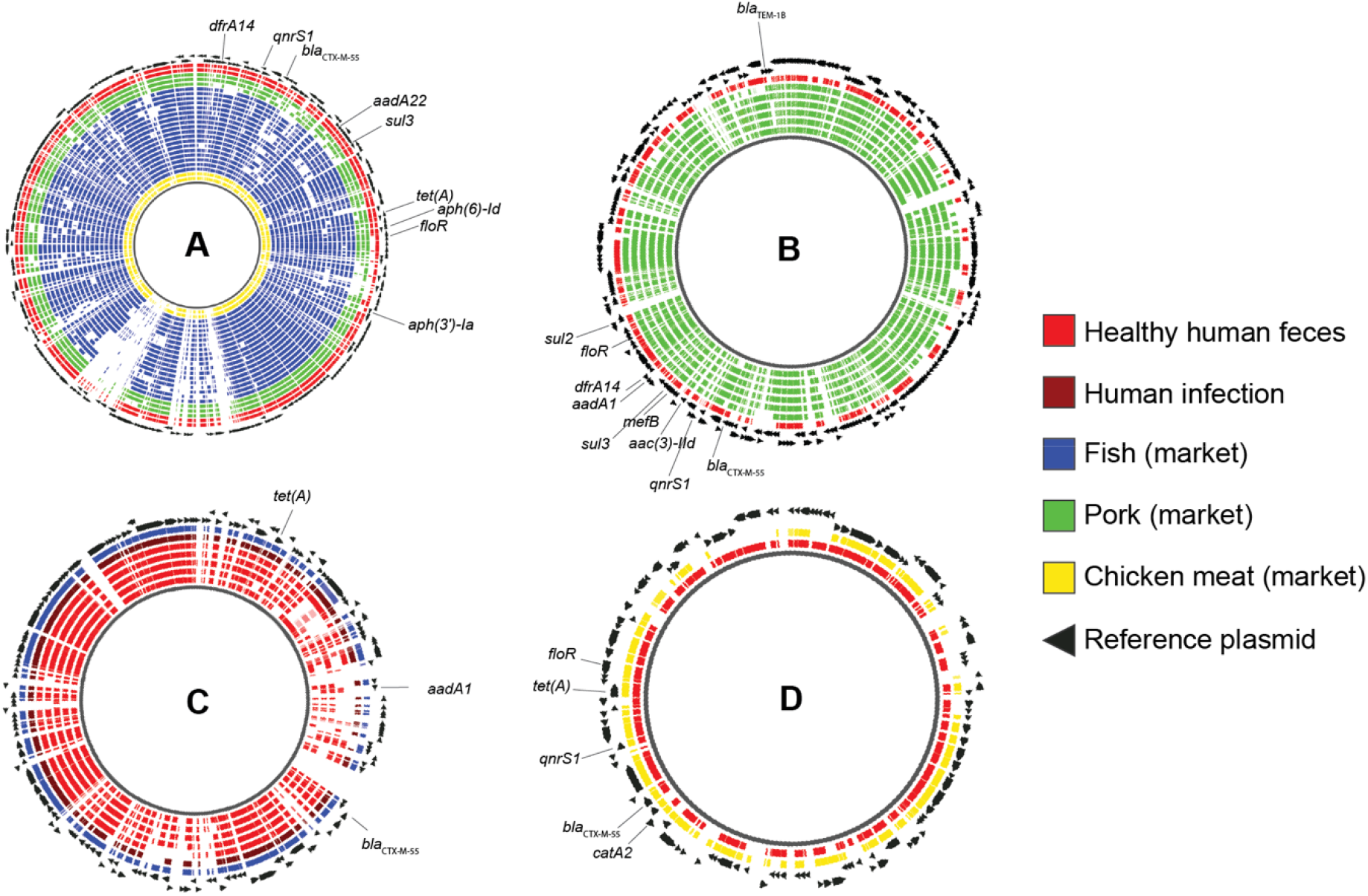
Occurrence of four *bla*_CTX-M-55_-encoding plasmids among *Escherichia coli* and *Salmonella enterica* from humans and food, Cambodia. A) pC27A-CTX-M-55, B) pP59A-CTX-M-55, C) pP225M-CTX-M-55, D) p276M-CTX-M-55. Plasmids A, B, and D harbored Tn*CTX-M/qnrS* (Fig. 4); plasmid C did not. *E. coli* and *S. enterica* strains harboring reference plasmids A through D were identified using the NASP pipeline; strains that mapped to at least 70% of the reference plasmid genome with at least 10x sequencing depth were considered matches. Regions of similarity between these matches and each reference plasmid were visualized using the GView server using default parameters (80% identity threshold for coding sequences). Each concentric circle represents a unique match and is colored by origin. The arrows indicate coding sequences identified on the reference plasmid.

In addition to being distributed among multiple vertebrate hosts, these four ESBL-encoding plasmids were also detected among *E. coli* from diverse genetic backgrounds (**Supplementary Text, Data S1**). This indicated that plasmid sharing across hosts was not exclusively driven by the zoonotic transmission of specific bacterial clones; rather, frequent mixing of host-adapted strains likely allowed for the uptake of each plasmid into diverse genetic contexts, a phenomenon others have observed in hospitals in high-income settings (Sheppard et al., 2016). Our findings suggest that in regions with high leakiness, focusing exclusively on the spillover of specific bacterial clones may fail to capture the spread of mobile resistance elements across a diversity of host-adapted strains. Frequent plasmid sharing could also create opportunities for the evolution of novel, antimicrobial-resistant human pathogens.

Next, we compared the four ESBL-encoding plasmid types to determine if the ESBL genes themselves shared a common origin. We identified a putative, *bla*_CTX-M_-encoding transposon that was integrated across three of the four plasmids (**Fig. 4**). This ∼6 kb transposon, hereafter referred to as Tn*CTX-M/qnrS*, also encoded *qnrS1* which confers low-level fluoroquinolone resistance. The *bla*_CTX-M-55_ and *bla*_CTX-M-15_ variants of the transposon, which differed by only one single nucleotide polymorphism, were highly prevalent among *Enterobacteriaceae*, including *E. coli, Klebsiella pneumoniae*, and *Salmonella enterica* from humans and animal products in Cambodia (**Fig. 2b, Fig. S2, Data S1**). Examination of a global *Escherichia/Shigella* database (https://enterobase.warwick.ac.uk/species/index/ecoli) revealed that the *bla*_CTX-M-15_ variant was broadly dispersed, while the *bla*_CTX-M-55_ variant was mainly identified in isolates from Asia including Vietnam, Laos, Thailand, Taiwan, and China (**Supplementary Text, Data S2**). The mobilization of the *bla*_CTX-M-55_ variant across different plasmids, bacterial lineages, and hosts strongly suggests that bacterial exchanges among humans and animals are exceptionally frequent in this region.

**Fig. 4.**
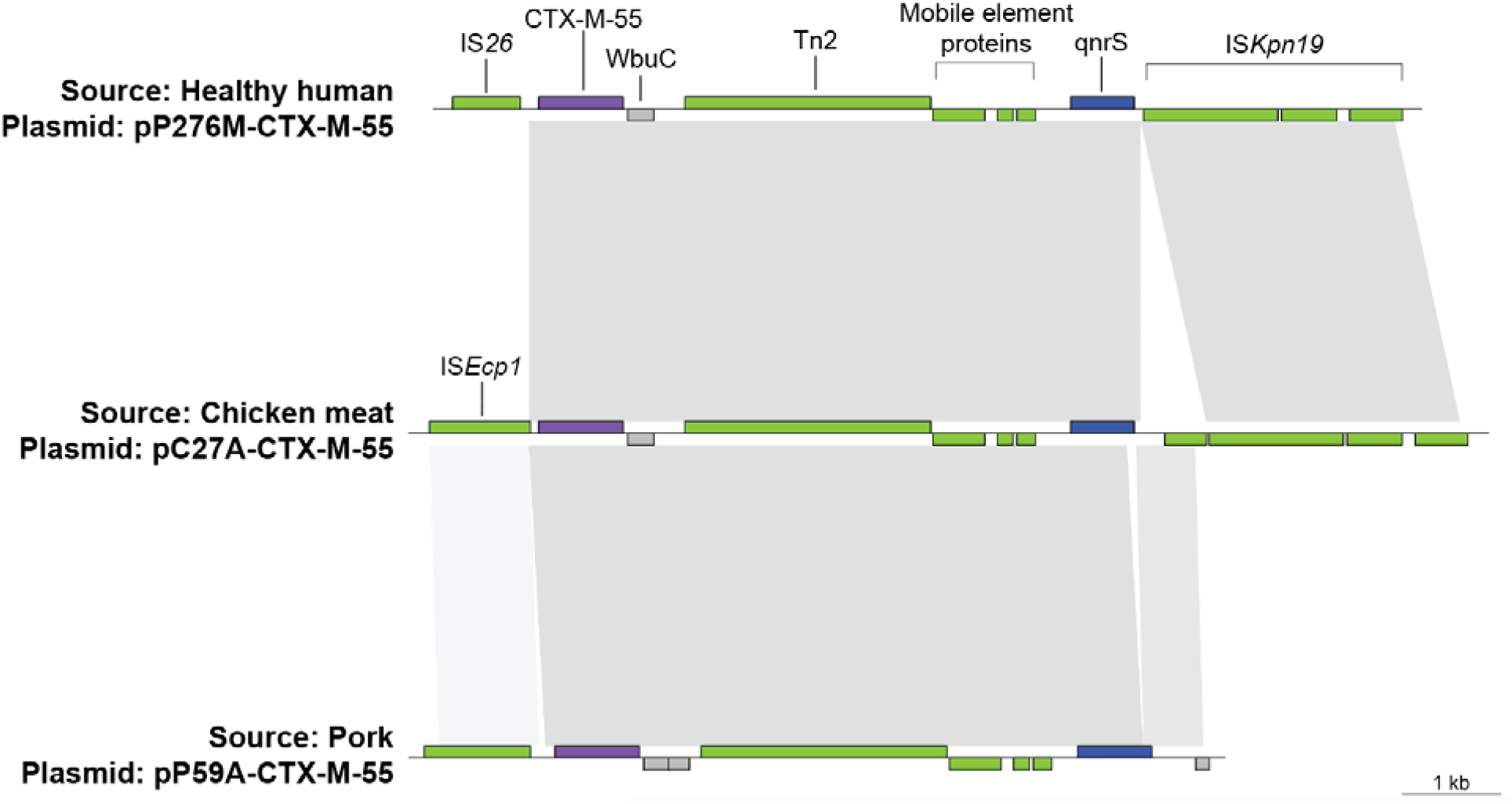
Encoded proteins in ∼6 kb putative transposon harboring *bla*_CTX-M-55_ and *qnrS1*, referred to as Tn*CTX-M/qnrS*, as identified in multiple plasmids of human and animal-origin *E. coli*. *<u>Note</u>*: CTX-M-55 is highlighted in purple, qnrS in blue, mobile element proteins in green, and all other proteins are gray. Shading indicates regions with >97% identity; darker gray indicates higher identity. CTX-M-15 and CTX-M-3-producing variants were observed in other isolates.

The pervasive leakiness between humans and animals in countries such as Cambodia may make it impossible to establish the host origins of mobile resistance elements. All three plasmids identified in this study that encoded Tn*CTX-M/qnrS* (**Fig. 4**) also carried the *floR* gene, which confers resistance to chloramphenicol. Given that phenicols are widely used in pigs and chickens but rarely in human medicine in Cambodia (Carrique-Mas et al., 2015; Choisy et al., 2019; G. Ström et al., 2018), this strongly supports the premise that these plasmids have been under selection in the food-animal production sector. Whether these plasmids acquired Tn*CTX-M/qnrS* before or after acquiring *floR*, and in which host, may be impossible to discern in this setting given the circular nature of bacterial exchange and the use of cephalosporins and fluoroquinolones in both humans and animals. Therefore, despite evidence of sector specific selection, expending resources to track the host origins of these plasmids may not be an effective approach for addressing antimicrobial resistance in such settings. Instead, cost-effective strategies for implementing environmental controls at the human-animal interface, *i*.*e*., “plugging the leaks,” should actively be explored.

This study was limited by our lack of access to samples collected directly from food animals, resulting in our use of meat and fish products as proxies. Theoretically, the bacteria that we isolated from these products could have originated from human contamination. However, 80% of the strains from meat and fish were resistant to phenicols (compared to <30% of the strains from humans) (Nadimpalli et al., 2019b), suggesting that food animals, which are regularly given phenicols (Carrique-Mas et al., 2015; Choisy et al., 2019; G. Ström et al., 2018), were the main source of contamination. A strength of this study was our use of long-read sequencing to establish the widespread dissemination of an ESBL-encoding transposon among *E. coli, S. enterica, K. pneumoniae*, and *Shigella* spp. recovered from multiple vertebrate hosts.

There is no doubt that antimicrobials are the greatest selective force for the emergence of new antimicrobial-resistant bacteria; however, poor environmental controls may be the greatest potentiators for such strains. In this study, detection of the same mobile resistance elements across multiple vertebrate hosts indicates that antibiotic stewardship alone may be insufficient to stem the growing problem of antimicrobial resistance in Southeast Asia. Instead, our findings indicate that antibiotic stewardship must be complemented by plugging the leaks between humans and food-animals. In addition to stemming the circular flow of antimicrobial-resistant bacteria, infrastructural WASH and biosecurity improvements will also reduce disease, and thereby, antibiotic demand, in both humans and animals.

Dietary changes in low and especially middle-income countries suggest the need to plug leaks is increasingly urgent. Specifically, as these countries develop, their demand for animal-based protein is soaring, driving up food-animal production and concomitant antimicrobial use (Van Boeckel et al., 2015). As a result, several countries that lack universal access to clean water and sanitation are expected to double their antimicrobial use in animal production over the next 10 years (Van Boeckel et al., 2015). Currently, less than 4% of funding through the Joint Programming Initiative for Antimicrobial Resistance (JPIAMR) is allocated towards identifying cost-effective environmental controls that curb microbial exchange (World Bank Group, 2019). This minimal investment may reflect a high-income country bias among scientists and policymakers who accept the generalizability of research conducted in settings with strong environmental controls, where the exchange of mobile resistance elements between food-animals and the broader community is limited (Day et al., 2019).

Our findings strongly suggest that the lack of environmental controls along with widespread antimicrobial use is leading to the dissemination of novel resistance elements and the evolution of antimicrobial-resistant human pathogens in Southeast Asia. Further studies are needed to determine the impacts of leakiness on the dissemination of antibiotic resistance in other settings, and additional research is needed to determine the most critical and most cost-effective leaks that can be plugged to prevent this. We propose that evaluating and implementing WASH and biosecurity initiatives should be prioritized along with antimicrobial stewardship to address the antimicrobial-resistance crisis in resource poor settings.

## Materials and Methods

### Whole genome sequencing and isolate selection

We previously isolated and characterized ESBL-producing *Escherichia coli* (ESBL-*Ec*) from gut-colonized, healthy humans (n=88) and market-origin chicken, pork, and fish (n=93) at the Institut Pasteur du Cambodge from 2015-2016 (Nadimpalli et al., 2019b). Briefly, food-origin third-generation cephalosporin-resistant (3GCR) *E. coli* were isolated from 60 specimens each of fish and pork and 30 specimens from chicken; human-origin 3GCR-*E. coli* were isolated from fecal swabs of recently pregnant healthy women living in the same neighborhood where markets were sampled. For one isolate per sample, ESBL production was confirmed via double-disk synergy testing and species identification by bioMérieux’s API 20E system. DNA extraction, library preparation, and whole genome sequencing were carried out on the Illumina NextSeq 500 platform using a 2×150 paired-end protocol (Nadimpalli et al., 2019b). The raw reads were pre-processed and assembled using SPAdes (Bankevich et al., 2012), assigned a multilocus sequence type (MLST) using the Achtman scheme (Larsen et al., 2012), and screened for acquired resistance genes using ResFinder v3.1.0 (selected threshold equal to 90% nucleotide identity) (Zankari et al., 2012).

As previously reported, we identified the *bla*_CTX-M-55_ gene to be the most common ESBL gene type among ESBL-*Ec* from food (62/93), and the second most common ESBL gene type among ESBL-*Ec* from humans (26/88). The *bla*_CTX-M-55_ gene was present across diverse sequence types among ESBL-*Ec* from food, but among healthy humans, *bla*_CTX-M-55_ was significantly more common among MLSTs belonging to clonal complex (CC) 10 (*p*<0.05 by two-sided Fisher’s exact test).

Thus, for the present study, we selected five *E. coli* CC10 isolates from food (n=3) and humans (n=2) for long-read sequencing using the MinION platform (Oxford Nanopore Technologies) to resolve the genetic context of their ESBL genes (**Fig. S1**).

### Long-read sequencing, hybrid genome assembly and genome characterization

DNA was extracted from 1 ml fresh overnight LB culture on blood agar with the GenFind v2 kit (Beckman Coulter) using a DynaMag-2 magnet (Thermo Fisher Scientific). A MinION sequencing library was prepared using the 1D Native Barcoding Sequencing Kit (SQK-LSK108 + EXP-NBD103) and sequenced in an R9.4.1 flow cell with a MinION Mk1B (Oxford Nanopore Technologies). Fast5 read files were base-called and demultiplexed with Albacore v2.3.0 (Oxford Nanopore Technologies). Sequencing adaptors were removed with Porechop v0.2.2 (Wick et al., 2017a) and reads quality filtered to at least q8 with NanoFilt v2.0.1 (De Coster et al., 2018) (assembly statistics are provided in **Table S1**). We used Unicycler v0.4.4 (Wick et al., 2017b) to construct high-quality hybrid assemblies using the Oxford Nanopore Technology long reads and pre-existing Illumina short-read sequences from the same isolates, trimmed with Trimmomatic v0.36 (Bolger et al., 2014) to remove sequencing adaptors and low quality (Qs20) ends. We used the CLC Genomics Workbench v10.1.1 (Qiagen) to manually correct the assemblies, which were then annotated with the Rapid Annotation Using System Technology Server (RAST) v2.0 (Aziz et al., 2008). We screened the assemblies for acquired antibiotic resistance genes using ResFinder v2.1 (Camacho et al., 2009) (selected threshold equal to 90% nucleotide identity). Plasmid replicons were detected using PlasmidFinder v2.0 (Carattoli et al., 2014) (selected threshold equal to 90% identity) and incompatibility group F (IncF) subtyping was performed using PubMLST (**Fig. S1**).

Draft genomes have been uploaded to NCBI’s GenBank under Bioproject number PRJNA566431. Individual accession numbers are available in **Table S2**.

### Identification of shared *bla*_CTX-M-55_-encoding region

We aligned the *bla*_CTX-M-55_-encoding contigs from each draft genome with ProgressiveMauve v2.4.0 (Darling et al., 2010) to identify if there was a shared genetic context that was smaller than a plasmid between strains (*e*.*g*. transposon) (**Fig. S1**). We further annotated the identified region using BLASTP and characterized insertion elements using ISFinder (Siguier et al., 2006).

### Detection of *bla*_CTX-M_-encoding elements among Cambodian ESBL-producing

### *Enterobacteriaceae* collections

We screened multiple ESBL-producing *Enterobacteriaceae* collections from Cambodia for the *bla*_CTX-M_-encoding elements identified in this study. These collections included the aforementioned collection of ESBL-producing *E. coli* from food and healthy humans (n=181) (Nadimpalli et al., 2019b), ESBL-producing *E. coli* from clinical specimens, including urine and blood (n=15) (Nadimpalli et al., 2019b), ESBL-producing *Salmonella enterica* from food (n=26) (Nadimpalli et al., 2019a), and ESBL-producing *Klebsiella pneumoniae* from food (n=8) (**Data S1**). All methods for isolating, characterizing, and sequencing the *E. coli* and *Salmonella* collections have been described above and/or previously been published in detail (Nadimpalli et al., 2019b, 2019a), *K. pneumoniae* were isolated from the same meat and fish samples as *E. coli* and *Salmonella*, following the same culturing methods used to isolate 3GCR-*E. coli* (Nadimpalli et al., 2019b). *K. pneumoniae* were confirmed by MALDI-TOF. Library preparation was performed using the Nextera XT V2 300-cycle Kit (Illumina) and whole genome sequencing was carried out on the NextSeq 500 platform (Illumina) using a 2×150 paired-end protocol to achieve a minimum sequencing depth of 50-fold. Sequence data have been deposited in the European Nucleotide Archive (ENA; https://www.ebi.ac.uk/ena) under project number PRJEB34504.

All collections were screened for a) the four *bla*_CTX-M-55_-encoding plasmids identified in the hybrid assemblies and b) a smaller, putative *bla*_CTX-M_-encoding transposon that was shared among three plasmids, referred to as Tn*CTX-M/qnrS* (**Fig. S1**). To screen for the a) *bla*_CTX-M-55_-encoding plasmids, we used the Northern Arizona SNP Pipeline (NASP) v1.0.0. (Sahl et al., 2016) Specifically, we aligned raw reads to each plasmid using the Burrows-Wheeler Aligner (BWA) and identified SNPs using GATK UnifiedGenotyper. Strains that mapped to at least 70% of the reference plasmid with at least 10x sequencing depth were considered to harbor the plasmid. Regions of similarity between these matches and each reference plasmid were visualized using the GView server using default parameters (80% identity threshold for coding sequences, https://server.gview.ca/). To screen for b) Tn*CTX-M/qnrS* we took a 2-step approach. First, we extracted the protein-coding segments of this region and built a BLAST database. We then generated *de novo* assemblies of the Cambodian ESBL-producing *E. coli* genomes (n=196) using SPAdes (Bankevich et al., 2012) and queried them for the protein-coding genes in the transposon using BLASTN. We analyzed BLAST results using a custom R script and arbitrarily included as a hit any contig where at least 50% of the proteins were present with 99% coverage and 95% identity and where at least genes encoding a transposase protein and a CTX-M variant were present. Second, we screened the Cambodian ESBL-producing *E. coli* collections (n=196) for Tn*CTX-M/qnrS* using NASP, as described above. Strains that mapped to at least 99% of the reference region with at least 10x sequencing depth were considered to harbor the region of interest. Findings from the two approaches were comparable. Thus, we proceeded to use only the NASP pipeline to screen the remaining Cambodian *Salmonella* (n=26) and *K. pneumoniae* (n=8) collections for the Tn*CTX-M/qnrS* element. We only report matches identified through the NASP pipeline here.

We constructed core genome phylogenies of our *E. coli* and *S. enterica* datasets to visualize the presence of a) each reference plasmid and b) Tn*CTX-M/qnrS* by isolate source and sequence type or serotype. For each tree, we selected a published reference genome that belonged to the same sequence type or serotype as other isolates in each dataset; *i*.*e*., CP011113 for *E. coli* (ST10) and CP010282 for *S. enterica* (serotype Newport). SNP alignments were obtained by running NASP as described above and removing recombined regions using Gubbins v2.3.4 (Croucher et al., 2015). We annotated the resulting phylogenetic trees using iTOL (Letunic and Bork, 2016).

### Detection of Tn*CTX-M/qnrS* among a global *Enterobacteriaceae* collection

We used BLASTN to query the entire Escherichia/Shigella database of EnteroBase (https://enterobase.warwick.ac.uk/species/index/ecoli, assessed in 2017, n=∼60,000) for the Tn*CTX-M/qnrS* element (**Fig. S1**). Strains that contained this region with 90% coverage and 90% identity were considered to harbor this element.

## Data availability

The accession numbers for the five *Escherichia coli* draft genomes presented in this study are available in NCBI’s GenBank under Bioproject number PRJNA566431. The accession numbers for the Illumina sequences generated from the ESBL-producing *Escherichia coli* (n=196), *Salmonella enterica* (n=26), and *Klebsiella pneumoniae* (n=8) isolates described in this study are available in the European Nucleotide Archive (ENA; https://www.ebi.ac.uk/ena) under accession numbers PRJEB25898, PRJEB27759, and PRJEB34504, respectively.

## Acknowledgments

This work was supported by the Dennis and Mireille Gillings Foundation, Pasteur Foundation U.S., MSD AVENIR, Monaco Department of International Cooperation, and Institut Pasteur. M.LN. was supported by NIH award KL2TR002545 and the Stuart B. Levy Center for Integrated Management of Antimicrobial Resistance at Tufts (Levy CIMAR), a collaboration of Tufts Medical Center and the Tufts University Office of the Vice Provost for Research (OVPR) Research and Scholarship Strategic Plan (RSSP). F.G. was supported by a grant from the Région Normandie. L.B.P was supported by the Wellcome Trust. The authors declare no competing interests.

## Supplementary Materials

### Supplementary Text

Table S1. Assembly statistics for draft *Escherichia coli* genomes constructed using Oxford Nanopore Technology (ONT) long reads and Illumina short reads.

Table S2. Accession numbers for hybrid draft genomes, deposited in NCBI GenBank Bioproject PRJNA566431.

Fig S1. Methods Overview. Flowchart describing draft genome assembly, characterization of *bla*_CTX-M_-encoding mobile genetic elements (MGEs), and screening of local and global *Enterobacteriaceae* datasets for these MGEs. Shaded region labeled “Prior Work” indicates previously reported work that informed sample selection for this study.

Fig S2. Detection of the Tn*CTX-M/qnr* transposon element and plasmid pC27A-CTX-M-55 (IncHI1-type) in ESBL-producing *Salmonella enterica* recovered from meat and fish, Cambodia. The same ESBL-encoding elements detected in *E. coli* from humans and meat (Fig. 2 of main text) were detected in *Salmonella enterica*, indicating that the mobilization of these elements may be common among *Enterobacteriaceae* species in this setting. Only the *bla*_CTX-M-55_ variant of Tn*CTX-M/qnrS* was identified among *S. enterica*. Data S1. Detection of *bla*_CTX-M_-encoding elements among ESBL-producing *Enterobacteriaceae* from Cambodia.

Data S2. Detection of Tn*CTX-M/qnrS* transposon element among *Escherichia* spp. & *Shigella* spp. genomes in EnteroBase.

