## Supplemental Text for "Leakiness at the human-animal interface in Southeast Asia and implications for the spread of antibiotic resistance"

**This PDF file includes:**

Supplementary Text

Figs. S1-S2

Table S1-S2

**Other Supplementary Materials for this manuscript include the following:**

Data S1: Detection of *bla*_CTX-M_-encoding elements among ESBL-producing *Enterobacteriaceae* from Cambodia

Data S2: Detection of Tn*CTX-M/qnrS* transposon element among *Escherichia* spp. & *Shigella* spp. genomes in EnteroBase

**Supplementary Text**

Characterization of *bla*_CTX-M-55_-encoding plasmids

The *bla*_CTX-M-55_ gene was plasmid-encoded for all four CTX-M-55-producing *E. coli* we completed hybrid assemblies for; the one CTX-M-14-producing *E. coli* we assembled harbored the *bla*_CTX-M-14_ gene on its chromosome (**Table S1**).

Two of the four *bla*_CTX-M-55_-harboring plasmids we detected were highly similar (≥90% coverage, ≥95% identity) to previously reported plasmids. Plasmid pC27A-CTX-M-55 (size: 264 kb), an IncHI1-type plasmid originally detected in an *E. coli* ST7369 isolate from chicken meat, was similar to plasmids identified in multiple *S. enterica* serovars, including: Agona from wild birds in Australia (CP048776.1), Muenster from an animal sample in China (CP045038.1), Goldcoast from a foodborne outbreak in Taiwan (CP039170.1, CP037959.1),^1^ an unknown serovar recovered from a duck in China (MN539018.1), and Kentucky (CP039440.1) and Newport (CP039436.1) from chicken samples in Vietnam. Plasmid pP225M-CTX-M-55 (size: 142 kb), an IncF[F18:A-:B1]-type originally detected in an *E. coli* ST10 isolate from the feces from a healthy human, was similar to plasmids identified in *E. coli* from wetland sediment (MN158989.1, MN158992.1) and *mcr1*-harboring *E. coli* from humans (KX276657.1, CP029748.1).

The other two *bla*_CTX-M-55_-harboring plasmids we detected were substantially less similar to previously published plasmids. Plasmid pP59A-CTX-M-55 (size: 253 kb), an IncHI2-type plasmid originally detected in an *E. coli* ST10 isolate from pork meat, shared between 78-82% coverage and >99% identity with plasmids identified in *E. coli* from horses (KF362122.2) and a slaughterhouse in China (CP053721.1), *Salmonella* spp. from meat in China (MG874042.1) and food workers in Japan (AP020333.1), and *mcr1*-harboring *K. pneumoniae* from Taiwan (MH733010.1). Plasmid pP276M-CTX-M-55 had an unknown Inc type and did not closely match any previously published plasmid; the closest matches (63-69% coverage, >99% identity) were plasmids identified in *E. coli* from the environment (CP010243.1) and goose farms (CP034590.1, CP034735.1) in China.

Distribution of *bla*_CTX-M-55_-encoding plasmids among local *Enterobacteriaceae* collections

The *bla*_CTX-M-55_-encoding plasmids we characterized were harbored by Cambodian-origin *E. coli* belonging to diverse sequence types (**Data S1**). Plasmid pC27A-CTX-M-55 (IncHI1-type) was identified among 18 different *E. coli* sequence types recovered from healthy human feces, chicken meat, and pork meat; no two sources were contaminated with pC27A-CTX-M-55-harboring *E. coli* belonging to the same sequence type. Plasmid pP225M-CTX-M-55 (IncF-type) was identified among six different *E. coli* sequence types recovered from healthy human feces, a urinary tract infection, and market-origin fish. The same sequence type (ST4456) was detected in both human feces and a human infection; these specimens were collected from two different individuals more than five months apart. Plasmid pP59A-CTX-M-55 (IncHI2-type) was detected among four different *E. coli* sequence types recovered from healthy human feces and pork meat; *E. coli* ST10 was detected in both feces and pork meat. Plasmid pP276M-CTX-M-55 (unknown Inc type) was exclusively detected among *E. coli* ST48 recovered from a healthy human and from chicken meat.

Plasmid pC27A-CTX-M-55 was also detected among diverse *Salmonella enterica* serotypes, including Wandsworth, Virchow, Bovismorbificans, Newport, and Saintpaul (**Fig. S2**). The three other CTX-M-55-producing plasmids we characterized were not detected among our Cambodian collections of ESBL-producing *S. enterica* (n=26) and none of the four plasmids were detected among ESBL-producing *K. pneumoniae* (n=8) (**Data S1**).

Distribution of Tn*CTX-M/qnrS* in Cambodian and global *Enterobacteriaceae* collections

Tn*CTX-M/qnrS* was widely prevalent among ESBL-*Ec* from healthy humans in Cambodia (42/88) and among food-origin ESBL-producing *E. coli* (54/93), *S. enterica* (19/26), and *K. pneumoniae* (1/8). Almost all strains harboring this element encoded either *bla*_CTX-M-15_ or *bla*_CTX-M-55_ (116/117); one strain from human feces encoded *bla*_CTX-M-3_. Tn*CTX-M/qnrS* was present across multiple *E. coli* STs and *Salmonella* serotypes, with no clear association to a single clonal group. However, it was not detected in ESBL-*E*c causing bloodstream (0/12) or upper urinary tract (0/2) infections in any Cambodian-origin isolates that we screened.

Of ~60,000 *Escherichia coli/Shigella* genomes uploaded to EnteroBase in 2017, we detected Tn*CTX-M/qnrS* (90% coverage and 90% identity thresholds) among 213 *E. coli* belonging to multiple sequence types and among 47 *Shigella* spp. of various species, including *S. sonnei, S. flexneri, S. boydii,* and *S. dysenteriae* (**Data S2**). Only strains encoding either *bla*_CTX-M-15_ or *bla*_CTX-M-55_ were observed to harbor this element. Interestingly, Tn*CTX-M/qnrS* was identified among both *E. coli* and *Shigella* isolated from human infections, including a urinary tract infection in Thailand (*Shigella flexneri*; SRR5660163; 2016); gallbladder (*E. coli* ST69; SRR4294847; 2016) and urinary tract infections (*E. coli* ST162; SRR5445922; 2016) in the United States, and diarrhea caused by Enteroinvasive *E. coli* (EIEC) (ST99; 2014) in the United Kingdom. Other isolates harboring the Tn*CTX-M/qnrS* element may have been isolated from infections but metadata were not available to confirm.

Due to sparse metadata, we were unable to determine if the presence of the Tn*CTX-M/qnrS* element was associated with specific geographic regions or hosts. However, the *bla*_CTX-M-55_ variant of the element was almost exclusively detected among isolates of Asian origin (30/35) in the EnteroBbase collection; the earliest reported isolate was detected in 2011. Other reports from Asia have described the *bla*_CTX-M-55_ variant of Tn*CTX-M/qnrS* in food, food animals, and infected patients ^2,3^, with the most recent report among *Salmonella* Goldcoast causing a prolonged gastroenteritis outbreak in Taiwan.^1^ Interestingly, a recent study in China also documented the transfer of a small mobile element co-harboring *bla*_CTX-M-55_ and *qnrS1* between *Salmonella* and *E. coli*, including transfer from the chromosome.^2^ However, genome sequence data were not available to compare the sequence of this region to the element characterized here.

**Table S1.** Assembly statistics for draft *Escherichia coli* genomes constructed using Oxford Nanopore Technology (ONT) long reads and Illumina short reads.

|  | Isolate | | | | |
| --- | --- | --- | --- | --- | --- |
|  | P225M | P276M | C27_A | P43_A | P59_A |
| Source | Healthy human feces | Healthy  human feces | Chicken meat | Pork | Pork |
| ESBL enzyme | CTX-M-55 | CTX-M-55 | CTX-M-55 | CTX-M-55 | CTX-M-14 |
| ONT long reads | | | | | |
| Mean read length (bp) | 15,862 | 15,047 | 11,878 | 10,793 | 13,304 |
| Mean read quality | 9.5 | 9.6 | 9.4 | 9.1 | 9.2 |
| Median read length (bp) | 10,710 | 10,372 | 7,438.5 | 3,144 | 5,781 |
| Median read quality | 9.7 | 9.7 | 9.6 | 8.9 | 9.1 |
| Number of reads (n) | 34,673 | 48,050 | 64,920 | 24,673 | 14,585 |
| Read length N50 (bp) | 24,503 | 23,498 | 21,882 | 34,083 | 35,782 |
| Total bases (n) | 549,969,269 | 723,010,863 | 771,145,811 | 266,283,006 | 194,031,111 |
| Illumina short reads | | | | | |
| Read length (bp) | 150 | 150 | 150 | 150 | 150 |
| Number of reads (n) | 1,724,525 | 1,573,833 | 1,158,935 | 1,662,327 | 1,409,176 |

**Table S2. Accession numbers for hybrid draft genomes, deposited in NCBI GenBank Bioproject PRJNA566431.**

| Isolate_ID | Description | BioSample | Accession Number |
| --- | --- | --- | --- |
| P43A | Chromosome | SAMN12789776 | CP044291 |
| P43A | pP43A-1 | SAMN12789776 | CP044292 |
| P276M | Chromosome | SAMN12789775 | CP044293 |
| P276M | pP276M-CTX-M-55 | SAMN12789775 | CP044294 |
| P225M | Chromosome | SAMN12789774 | CP044346 |
| P225M | pP225M-CTX-M-55 | SAMN12789774 | CP044347 |
| P225M | pP225M-2 | SAMN12789774 | CP044348 |
| P59A | Chromosome | SAMN12789773 | CP044298 |
| P59A | pP59A-CTX-M-55 | SAMN12789773 | CP044299 |
| P59A | pP59A-2 | SAMN12789773 | CP044300 |
| P59A | pP59A-3 | SAMN12789773 | CP044301 |
| P59A | pP59A-4 | SAMN12789773 | CP044302 |
| P59A | pP59A-5 | SAMN12789773 | CP044303 |
| P59A | pP59A-6 | SAMN12789773 | CP044304 |
| C27A | Chromosome | SAMN12789772 | CP044305 |
| C27A | pC27A-CTX-M-55 | SAMN12789772 | CP044306 |
| C27A | pC27A-2 | SAMN12789772 | CP044307 |
| C27A | pC27A-3 | SAMN12789772 | CP044308 |
| C27A | pC27A-4 | SAMN12789772 | CP044309 |

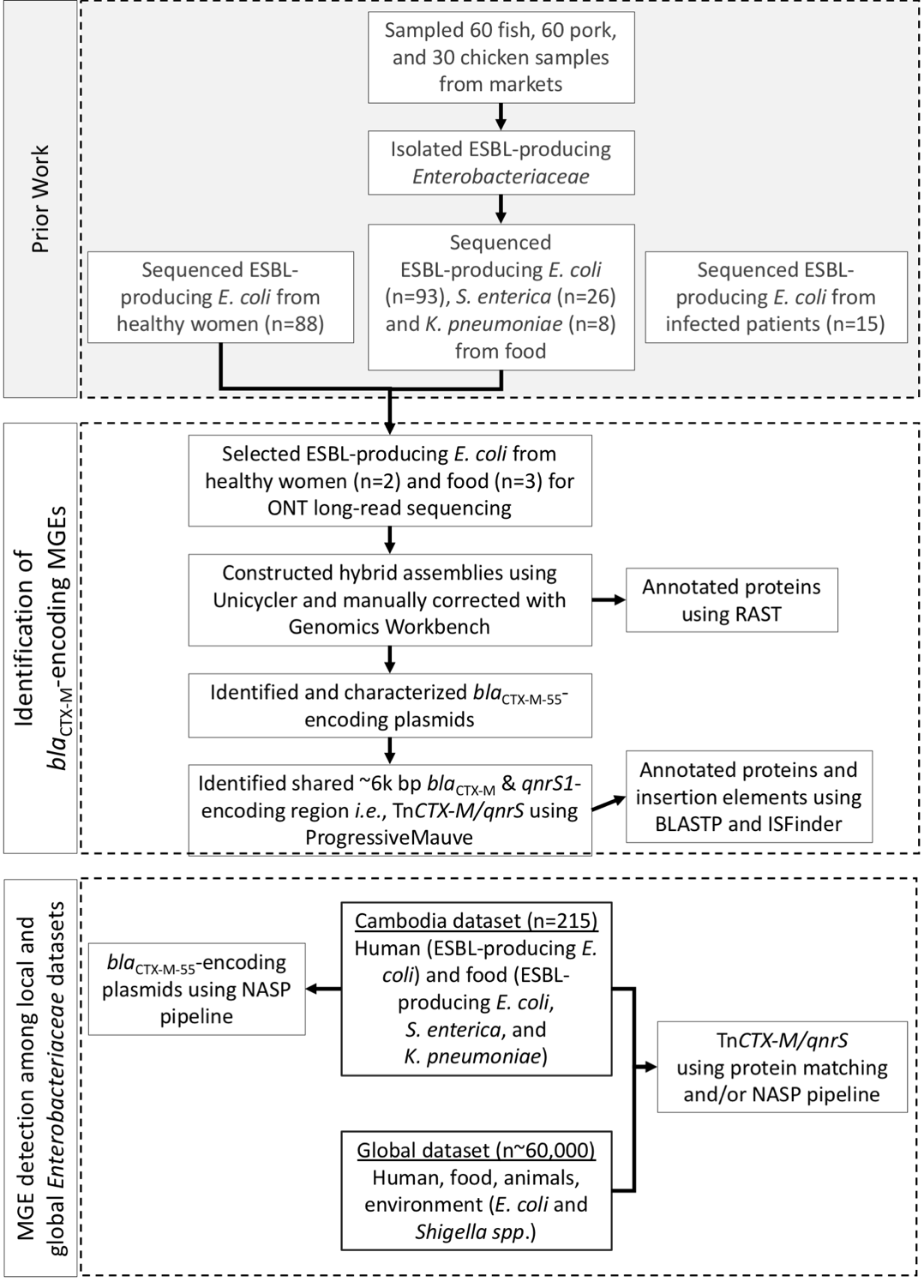

**Fig. S1. Methods Overview.** Flowchart describing draft genome assembly, characterization of *bla*_CTX-M_-encoding mobile genetic elements (MGEs), and screening of local and global *Enterobacteriaceae* datasets for these MGEs. Shaded region labeled "Prior Work" indicates previously reported work^4,5^ that informed sample selection for this study.

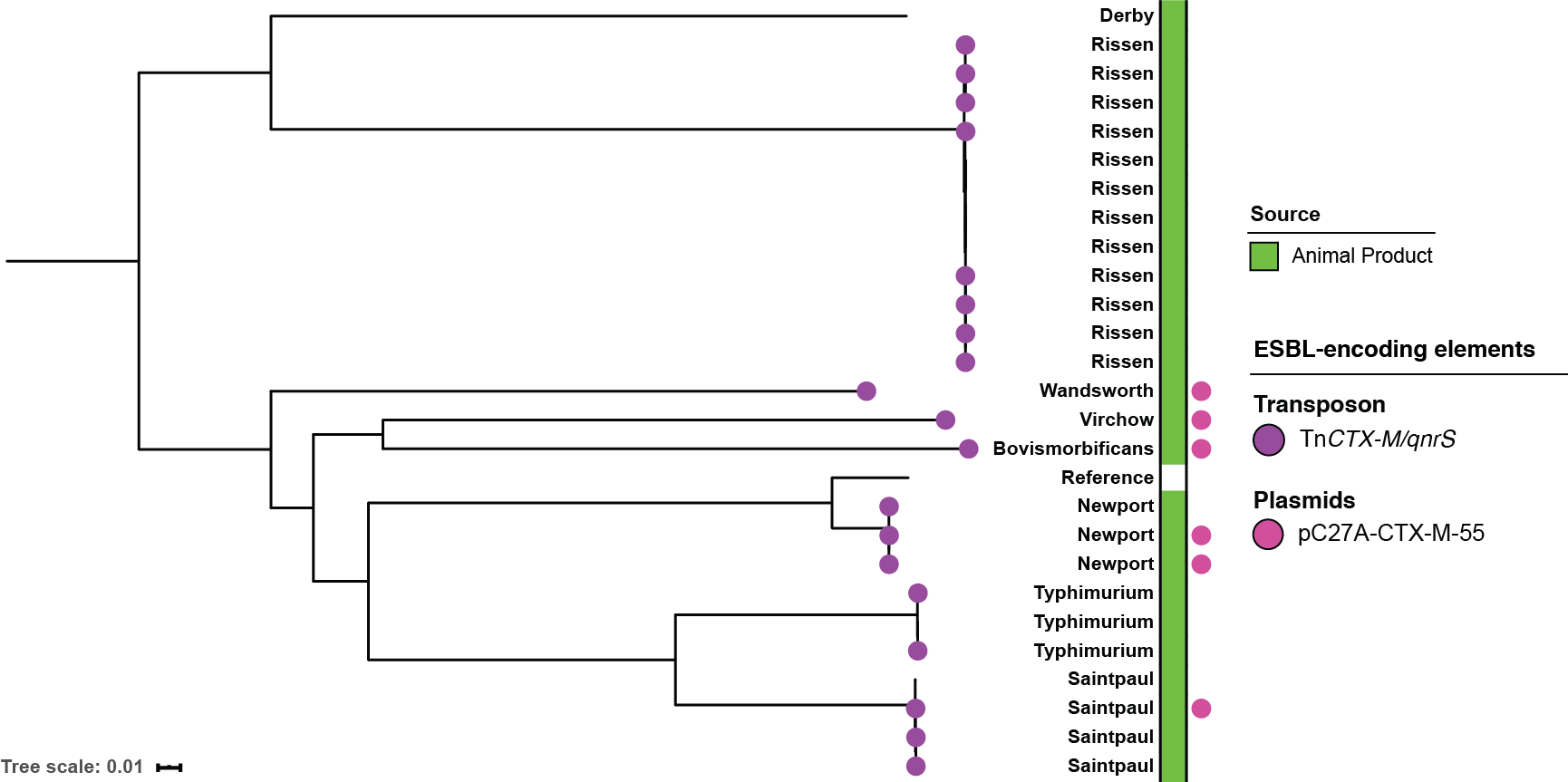

**Fig. S2. Detection of the Tn*CTX-M/qnr* transposon element and plasmid pC27A-CTX-M-55 (IncHI1-type) in ESBL-producing *Salmonella enterica* recovered from meat and fish, Cambodia**. The same ESBL-encoding elements detected in *E. coli* from humans and meat (Fig. 2 of main text) were detected in *Salmonella enterica*, indicating that the mobilization of these elements may be common among *Enterobacteriaceae* species in this setting. Only the *bla*_CTX-M-55_ variant of Tn*CTX-M/qnrS* was identified among *S. enterica*.
